# Altered sensory neuron development in CMT2D mice is site-specific and linked to increased GlyRS levels

**DOI:** 10.1101/2020.04.27.025098

**Authors:** James N. Sleigh, Aleksandra M. Mech, Tahmina Aktar, Yuxin Zhang, Giampietro Schiavo

## Abstract

Dominant, missense mutations in the widely and constitutively expressed *GARS1* gene cause a peripheral neuropathy that usually begins in adolescence and principally impacts the upper limbs. Caused by a toxic gain-of-function in the encoded glycyl-tRNA synthetase (GlyRS) enzyme, the neuropathology appears to be independent of the canonical role of GlyRS in aminoacylation. Patients display progressive, life-long weakness and wasting of muscles in hands followed by feet, with frequently associated deficits in sensation. When dysfunction is observed in motor and sensory nerves, there is a diagnosis of Charcot-Marie-Tooth disease type 2D (CMT2D), or distal hereditary motor neuropathy type V if the symptoms are purely motor. The cause of this varied sensory involvement remains unresolved, as are the pathomechanisms underlying the selective neurodegeneration characteristic of the disease. We have previously identified in CMT2D mice that neuropathy-causing *Gars* mutations perturb sensory neuron fate and permit mutant GlyRS to aberrantly interact with neurotrophin receptors (Trks). Here, we extend this work by interrogating further the anatomy and function of the CMT2D sensory nervous system in mutant *Gars* mice, obtaining several key results: 1) sensory pathology is restricted to neurons innervating the hindlimbs; 2) perturbation of sensory development is not common in mouse models of neuromuscular disease; 3) *in vitro* axonal transport of signalling endosomes is not impaired in afferent neurons of all CMT2D mouse models; and 4) *Gars* expression is selectively elevated in a subset of sensory neurons and linked to sensory developmental defects. These findings highlight the importance of comparative neurological assessment in mouse models of disease and shed light on key proposed neuropathogenic mechanisms in *GARS1*-linked neuropathy.

## 1. Introduction

Characterised by distal dysfunction of motor and sensory nerves, Charcot-Marie-Tooth disease (CMT) is a hereditary peripheral neuropathy that usually presents in adolescence and affects 1 in 2,500-5,000 people, which makes it the most common inherited neuromuscular condition (Pipis *et al.*, 2019). Classically, the disease can be categorised as CMT1, typified by demyelination and thus reduced nerve conduction velocity, CMT2 in which there is axon loss but no diminished nerve conduction velocity, and intermediate CMT that shares features of both CMT1 and CMT2 (Reilly *et al.*, 2011). Consistent with length-dependency, patients display slowly progressive, bi-lateral muscle weakness and sensory deficits predominantly in the extremities, typically starting in the feet. CMT is on a phenotypic spectrum with distal hereditary motor neuropathy (dHMN) and hereditary sensory/autonomic neuropathy (HSN/HSAN), which have mainly motor and sensory/autonomic involvement, respectively, and can be caused by mutations in the same gene (Pisciotta and Shy, 2018).

To date, mutations in more than 100 different genetic loci have been linked to CMT (Rossor et al., 2013; Pipis et al., 2019). Many causative CMT1 genes are selectively expressed by myelinating Schwann cells or have myelin-specific functions, providing mechanistic justification for pathology. However, CMT2-associated genes are involved in a variety of processes critical to general cell viability (e.g. mitochondrial dynamics, endolysosomal sorting, ubiquitination, heat shock response), and the pathomechanisms underlying neuronal selectivity remain relatively obscure. Fitting with this, the widely and constitutively active aminoacyl-tRNA synthetase (ARS) enzymes, which covalently bind specific amino acids to their partner tRNAs for protein translation (Ibba and Soll, 2000), represent the largest protein family implicated in CMT aetiology. To date, dominant mutations in five ARS-encoding genes (*GARS1*, *YARS1*, *AARS1*, *HARS1* and *WARS1*) have been identified to cause CMT (Wei *et al.*, 2019).

Encoding glycyl-tRNA synthetase (GlyRS), which charges glycine, *GARS1* is the first and best studied ARS gene linked to CMT (designated CMT type 2D, CMT2D, OMIM: 601472) (Antonellis *et al.*, 2003). Uncharacteristically and contravening length-dependency, CMT2D patients frequently display upper limb predominance with weakness beginning in dorsal interosseus muscles of the hand and progressing to involve lower limbs in about only half of patients (Sivakumar *et al.*, 2005; Antonellis *et al.*, 2018). Genetic studies across yeast, *Drosophila melanogaster* and mouse models for CMT2D indicate that, although neuropathy-causing mutations can abolish canonical GlyRS function and loss-of-function pathogenesis hypotheses prevail (Meyer-Schuman and Antonellis, 2017), the disease is most likely caused by a toxic gain-of-function (Boczonadi *et al.*, 2018; Wei *et al.*, 2019). Commensurate with mutant protein toxicity, wild-type *GARS1* overexpression in CMT2D mice has no discernible rescue effect on neuromuscular pathologies, while increased dosage of disease-causing *Gars* alleles causes more severe neuropathy (Motley *et al.*, 2011). Moreover, all assessed GlyRS mutants possess a similar conformational opening that excavates neomorphic surfaces usually buried within the structure of the wild-type enzyme (He *et al.*, 2011, 2015). Given that GlyRS is secreted from several different cell types (Park *et al.*, 2012; Grice *et al.*, 2015; He *et al.*, 2015; Park *et al.*, 2018), these uncovered protein regions can mediate aberrant deleterious interactions both inside and outside the cell (He *et al.*, 2015; Sleigh *et al.*, 2017a; Mo *et al.*, 2018), likely accounting for non-cell autonomous aspects of pathology (Grice *et al.*, 2015, 2018). While some of these mis-interactions are with neuronally-enriched proteins, the pathomechanisms underlying neuronal selectivity in CMT2D remain unresolved. Nevertheless, several recent studies indicate that impairments in the processes of axonal transport (Benoy *et al.*, 2018; Mo *et al.*, 2018) and protein translation (Niehues *et al.*, 2015) may be playing a causative role.

Several different mouse models are available for CMT2D (Seburn *et al.*, 2006; Achilli *et al.*, 2009; Morelli *et al.*, 2019), which have mutations in endogenous mouse *Gars*, causing phenotypes akin to human neuropathy. These mice display loss of lower motor neuron connectivity and disturbed neurotransmission at the neuromuscular junction (NMJ), causing muscle weakness and motor function deficits (Sleigh *et al.*, 2014a; Spaulding *et al.*, 2016). Furthermore, there appears to be a pre-natal perturbation of sensory neuron fate in dorsal root ganglia (DRG), such that CMT2D mice have more nociceptive (noxious stimulus-sensing) neurons and fewer mechanosensitive (touch-sensing) and proprioceptive (body position-sensing) neurons (Sleigh *et al.*, 2017a). Perhaps causing this and providing rationale for neuronal selectivity, mutant GlyRS mis-interacts with the extracellular region of tropomyosin receptor kinase (Trk) receptors. These largely neuron-specific transmembrane proteins mediate the development and survival of sensory neurons by binding with differential affinity to neurotrophins secreted from distal target cells/tissues (e.g. Schwann cells and muscles) (Huang and Reichardt, 2003). Activated neurotrophin-Trk receptor complexes are internalised in the periphery, sorted into signalling endosomes, and then retrogradely transported along microtubules to neuronal somas, where they elicit transcriptional events fundamental to nerve survival (Villarroel-Campos *et al.*, 2018).

The earliest manifestation of CMT2D in many individuals is transient cramping and pain in the hands upon cold exposure (Antonellis *et al.*, 2018). In addition to muscle weakness, this is followed by compromised reflexes and loss of sensation to vibration, touch, temperature and pin-prick (Sivakumar *et al.*, 2005). These symptoms are well replicated in *Gars*-neuropathy mice, highlighting their potential for studying sensory pathomechanisms (Sleigh *et al.*, 2017a).

However, the motor symptoms of CMT2D patients are often the focus of clinical investigation, given their relative severity. Moreover, *GARS1* neuropathy patients can show little to no sensory involvement and are thus diagnosed with dHMN type V (OMIM 600794) (Antonellis *et al.*, 2003). The pathological impact of mutant GlyRS on the sensory nervous system is therefore relatively under-studied and requires further attention if we are to elucidate the cause of its varied involvement in *GARS1*-linked neuropathy. Here, we have thus extended our sensory analyses in CMT2D mice to better understand the importance of anatomical location to pathology and to assess the relevance of some proposed disease mechanisms in afferent nerves.

## 2. Materials and Methods

### 2.1 Animals

All experiments were carried out following the guidelines of the UCL Queen Square Institute of Neurology Genetic Manipulation and Ethics Committees and in accordance with the European Community Council Directive of 24 November 1986 (86/609/EEC). *Gars*^*C201R*/+^(RRID: MGI 3849420) and SOD1^G93A^ (RRID: IMSR_JAX 002726) mouse handling and experiments were carried out under license from the UK Home Office in accordance with the Animals (Scientific Procedures) Act 1986 and were approved by the UCL Queen Square Institute of Neurology Ethical Review Committee. *Gars*^*Nmf249*/+^ (RRID: MGI 5308205) tissue was provided by Drs. Emily Spaulding and Robert Burgess (The Jackson Laboratory, Bar Harbor, ME), as previously described (Sleigh *et al.*, 2017a). *Gars*^*C201R*/+^ and *Gars*^*Nmf249*/+^ mice were maintained as heterozygote breeding pairs on a C57BL/6J background and genotyped as previously described (Seburn *et al.*, 2006; Achilli *et al.*, 2009). Both males and females were used in the analyses of mutant *Gars* alleles, as no clear sex-specific differences have yet been observed or reported. Genotyped using standard procedures (Gurney *et al.*, 1994), transgenic male mice heterozygous for the mutant human *SOD1* gene (*G93A*) on a mixed C57BL/6 SJL background (B6SJLTg [SOD1*G93A]1Gur/J) and wild type male littermate controls were used for the SOD1^G93A^ experiments. *Gars*^*C201R*/+^ mice sacrificed for one month and three month timepoints were 29-37 and 89-97 days old, respectively. The *Gars^Nmf249/+^* mice used at one month were P31-32, while SOD1^G93A^ mice were P30-31 and P100-101.

### 2.2 Tissue dissection

DRG were extracted from either non-perfused or saline-perfused mice as previously described (Sleigh *et al.*, 2016). The most caudal pair of floating ribs and the large size of lumbar level 4 (L4) DRG and associated axon bundles were used as markers to consistently and accurately define the spinal level. The forepaws of embryonic day 13.5 (E13.5) embryos were removed between the wrist and elbow joints, as outlined elsewhere (Wickramasinghe *et al.*, 2008).

### 2.3 Tissue immunofluorescence

Dissected DRG and E13.5 forepaws were processed for immunofluorescence and analysed as previously described in detail (Sleigh *et al.*, 2017a). The following antibodies were used: rabbit anti-GlyRS (1/200, Abcam, ab42905, RRID: AB_732519), rabbit anti-LysRS (1/200, Abcam, ab129080, RRID: AB_11155689), mouse anti-neurofilament (1/50, 2H3, developed and deposited by Jessell, T.M. / Dodd, J., Developmental Studies Hybridoma Bank, supernatant), mouse anti-NF200 (1/500, Sigma, N0142, RRID: AB_477257) and rabbit anti-peripherin (1/500, Merck Millipore, AB1530, RRID: AB_90725). The analyses of L1-L5 and C4-C8 wild-type DRG were performed at different times, as were the assessments of lumbar DRG dissected from the different genetic strains.

### 2.4 Western blotting of DRG lysates

Probing and analysis of DRG lysate western blots was performed as previously described (Sleigh *et al.*, 2017a), using the following antibodies: mouse anti-Gapdh (1/3,000, Merck Millipore, MAB374, RRID: AB_2107445), anti-GlyRS (1/2,000), anti-LysRS (1/500), anti-NF200 (1/1,000), anti-peripherin (1/1,000), rabbit anti-TrkB (1/1,000, BD Biosciences, 610101, RRID: AB_397507) and rabbit anti-TyrRS (1/500, Abcam, ab150429, RRID: AB_2744675). 10 μg of DRG lysate was loaded per lane.

### 2.5 Culturing primary DRG neurons

20-24 lumbar to thoracic DRG (Figure 1B) were dissociated and cultured on 35 mm glass bottom dishes (MatTek, P35G-1.5-14-C) in the presence of freshly added 20 ng/ml mouse glial cell line-derived neurotrophic factor (GDNF, PeproTech, 450-44) as detailed elsewhere (Sleigh *et al.*, 2017a). To reduce variability, a wild-type and *Gars*^*C201R*/+^ littermate of the same sex were dissected and cultured in parallel for each experimental replicate.

**Figure 1.**
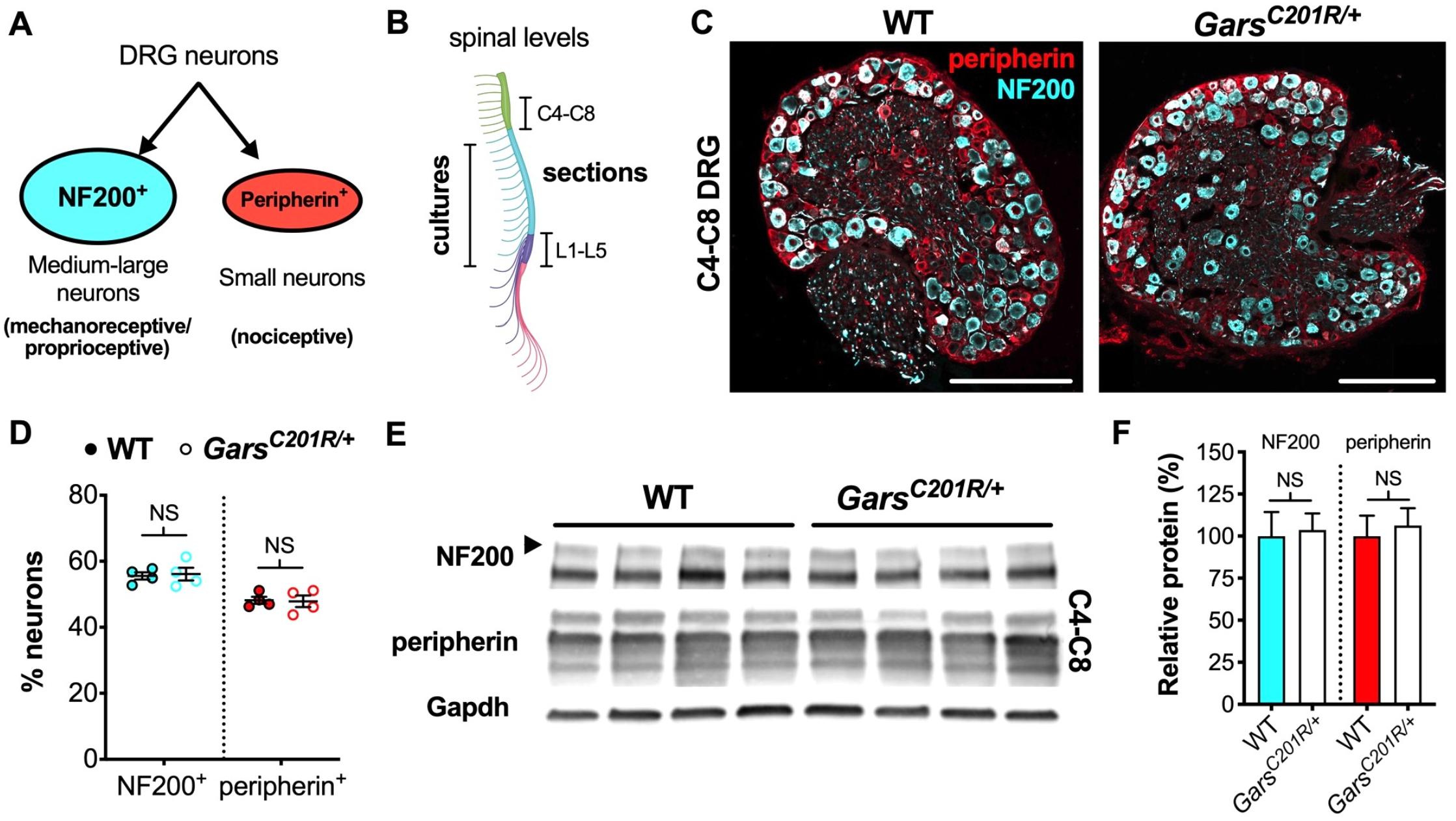
Sensory neuron development is not impaired in *Gars*^*C201R*/+^ DRG at cervical spinal levels. (**A**) Neurons found in sensory ganglia can be classified into NF200^+^ cells, which are mainly medium-large in size and function as either mechanoreceptors or proprioceptors, and peripherin^+^ cells that are generally small and nociceptive. (**B**) DRG used in this study were taken from cervical spinal level 4 (C4) to C8 and lumbar level 1 (L1) to L5 for immunofluorescence analysis and from thoracic to lumbar levels for primary cultures. The schematic was created with BioRender (https://biorender.com). (**C**) Representative immunofluorescence images of one month old wild-type and *Gars*^*C201R*/+^ cervical DRG sections stained for NF200 (cyan) and peripherin (red). Scale bars = 200 μm. (**D**) There was no difference in the proportions of NF200^+^ (*P* = 0.812, unpaired *t*-test) or peripherin^+^ (*P* = 0.885, unpaired *t*-test) neurons between genotypes in C4-C8 ganglia. *n* = 4. (**E**) Representative western blot of C4-C8 DRG lysates from one month old wild-type and *Gars*^*C201R*/+^ mice probed for NF200, peripherin, and the loading control Gapdh. (**F**) Consistent with the immunofluorescence analysis (D), there was no difference between genotypes in levels of NF200 (*P* = 0.835, unpaired *t*-test) or peripherin (*P* = 0.702, unpaired *t*-test) protein. *n* = 5. *NS*, not significant; *WT*, wild-type. See also **Supplementary Figure S1** and **S2**.

### 2.6 *In vitro* signalling endosome transport assay

The atoxic binding fragment of tetanus neurotoxin (H_C_T) was bacterially expressed and labelled with AlexaFluor647 antibody labelling kit (Life Technologies, A-20186) as previously outlined (Gibbs *et al.*, 2016). 24 h post-plating of dissociated DRG neurons, H_C_T-647 was added to the neuronal media at a final concentration of approximately 5 pg/ml, before gentle mixing by rotation and returning to 37°C for 25 min. H_C_T-containing medium was then aspirated, the cells were washed with 2 ml pre-warmed medium, before being slowly flooded with 2 ml standard medium containing all supplements. Within 10-90 min of the media change, endosome transport was imaged on an inverted LSM780 laser scanning microscope (Zeiss) inside an environmental chamber pre-warmed and set throughout the experiment to 37°C. An area containing a single neuronal process retrogradely transporting fluorescent endosomes was imaged using a 63x Plan-Apochromat oil immersion objective (Zeiss). Images were taken at 100x digital zoom (1024×1024, 1% laser power) every 2.4 s on average. Before selecting a neuronal process for analysis, it was first traced back to the cell body to confirm directionality of transport and imaged for area measurement (see below). Cultures from wild-type and *Gars*^*C201R*/+^ mice were imaged in the same session and, to avoid introducing time-dependent biases, their order was alternated across replicates. Two males and two females of each genotype were analysed at each timepoint.

### 2.7 Endosome transport analysis

Individual endosomes were manually tracked using Tracker (Kinetic Imaging Ltd.) as described previously (Sleigh *et al.*, 2020). Briefly, endosomes were included in the analysis if they could be observed for five consecutive frames and did not pause for >10 consecutive frames. Endosomes moving solely in the anterograde direction were infrequent and not included in the analysis. Individual frame-to-frame step speeds are included in the frequency histogram (an average of 2,370 ± 135 frame-to-frame speeds per animal), meaning that an endosome tracked across 21 consecutive frames will generate 20 frame-to-frame speeds in this graph. To determine the endosome speed per animal, individual endosome speeds were calculated and then the mean of these determined (an average of 95.1 ± 3.5 endosomes per animal). All speed analyses include frames and time during which endosomes may have been paused, *i.e.* the speed across the entire tracked run length is reported and not the speed solely when motile. An endosome was considered to have paused if it remained in the same position for two or more consecutive frames. The ‘% time paused’ is a calculation of the length of time all tracked endosomes remained stationary, while the ‘% pausing endosomes’ details the proportion of endosomes that displayed at least one pause while being tracked. An average of 26.2 endosomes were tracked per neuron, and at least three individual neurons were assessed per animal replicate.

### 2.8 Image analysis

Cell body areas of neurons analysed in endosome transport assays were measured using the freehand tool on ImageJ (https://imagej.nih.gov/ij/) to draw around the circumference of the somas. The diameters of neuronal processes imaged for transport were measured in ImageJ using the straight line tool. The average of five measurements across the width of the process was calculated. To determine in which cells GlyRS levels were highest in *Gars*^*Nmf249*/+^ lumbar DRG sections, all cells with increased GlyRS expression were first identified by eye in the single fluorescence channel. Cells positive for NF200 were then independently designated in the second channel. The percentage of GlyRS-elevated cells also positive for N200 was then calculated. Similarly, the percentage of NF200^+^ cells without an increase in GlyRS was also determined. All sections used for GlyRS analysis were stained and imaged in parallel with the same confocal settings to permit side-by-side comparison.

### 2.9 Statistical analysis

Data were assumed to be normally distributed unless evidence to the contrary could be provided by the D’Agostino and Pearson omnibus normality test. GraphPad Prism 8 (version 8.4.0, La Jolla, CA) was used for all statistical analyses. Means ± standard error of the mean are plotted, as well as individual data points in all graphs except for those depicting western blot densitometry. Unpaired *t*-tests and two-way ANOVAs were used throughout the study. Rather than ANOVAs, unpaired *t*-tests were used to analyse the percentages of NF200^+^ and peripherin^+^ neurons separately, because the two markers are not independently expressed. Similarly, western blot densitometry was also analysed using unpaired *t*-tests; since expression was calculated relative to wild-type levels for each individual protein, the expression of proteins in wild-type animals are not statistically comparable.

## 3. Results

### 3.1 Altered sensory development occurs specifically in lumbar segments of CMT2D mice

We previously showed that sensory neuron fate is altered during development in the mild *Gars*^*C201R*/+^ and more severe *Gars*^*Nmf249*/+^ mouse models for CMT2D, the extent of which correlated with overall model severity (Sleigh *et al.*, 2017a). By co-staining DRG for NF200, a marker of medium-large area mechanosensitive/proprioceptive neurons, and peripherin, which identifies small area nociceptive neurons (Figure 1A), we determined that mutant *Gars* DRG had fewer touch-and body position-sensing (NF200^+^) neurons and a concomitant increase in noxious stimulus-sensing (peripherin^+^) neurons. This phenotype was present at birth and did not change up to three months of age, suggesting it is developmental in origin and non-progressive. Ganglia assessed in these original experiments were isolated from lumbar level 1 (L1) to L5, which contains neurons that innervate the lower leg (Mohan *et al.*, 2014); however, CMT2D patients frequently display upper limb predominance (Antonellis *et al.*, 2018).

To determine whether the phenotype is also observed in forelimb-innervating ganglia (Tosolini *et al.*, 2013), we isolated and immunohistochemically analysed cervical level 4 (C4) to C8 DRG from one month old wild-type and *Gars*^*C201R*/+^ mice (**Figure 1B**). Co-labelling DRG for NF200 and peripherin (**Figure 1C**) and calculating the percentages of neurons expressing each marker, we saw no difference between genotypes (**Figure 1D**). Corroborating this, western blotting of cervical DRG lysates showed no difference in NF200 or peripherin protein levels (**Figure 1E** and **F**). Together, these data indicate that there is no impairment in sensory neuron identity in C4-C8 ganglia of *Gars*^*C201R*/+^ mice.

To confirm and extend the lumbar phenotype, we assessed levels of the protein TrkB, which binds brain-derived neurotrophic factor (BDNF) and neurotrophin-4 (NT-4) to ensure the survival of a mechanosensitive sub-population of NF200^+^ neurons (Montaño *et al.*, 2010). We found that lumbar ganglia of *Gars*^*C201R*/+^ mice have less full-length TrkB, consistent with there being fewer NF200^+^ neurons in the mutant DRG (**Supplementary Figure S1**).

We then statistically compared the proportions of wild-type cervical DRG neurons with previously published data from one month old wild-type L1-L5 DRG (NF200^+^ 40.7 ± 1.9%; peripherin^+^ 61.5 ± 2.1%) (Sleigh *et al.*, 2017a). We found that the ratio of subtypes is more even in cervical ganglia, which possess significantly more NF200^+^ and significantly fewer peripherin^+^ neurons than lumbar DRG (**Supplementary Figure S2A**).

In the past, we also identified a sensory neurodevelopmental phenotype in embryonic *Gars*^*C201R*/+^ hindlimbs (Sleigh *et al.*, 2017a). Dissecting hindpaws from embryonic day 13.5 (E13.5) mice and staining neurons for neurofilament (2H3), we observed impaired arborisation of nociceptive neurons found in the developing dorsal floor plate. To evaluate whether, this phenotype is also seen in forelimbs, we analysed forepaws from E13.5 embryos (**Figure 2A**). Similar to the hindpaws, there was no difference in sensory nerve growth between genotypes, assessed by measuring the distance from nerve growth cone to digit tip (**Figure 2B**). However, CMT2D forepaws did not display the nociceptive nerve branching defect present in lower limbs (**Figure 2C**). Therefore, similar to the DRG, developing sensory neurons originating at cervical spinal levels do not show the impairments found in lumbar afferent nerves.

**Figure 2.**
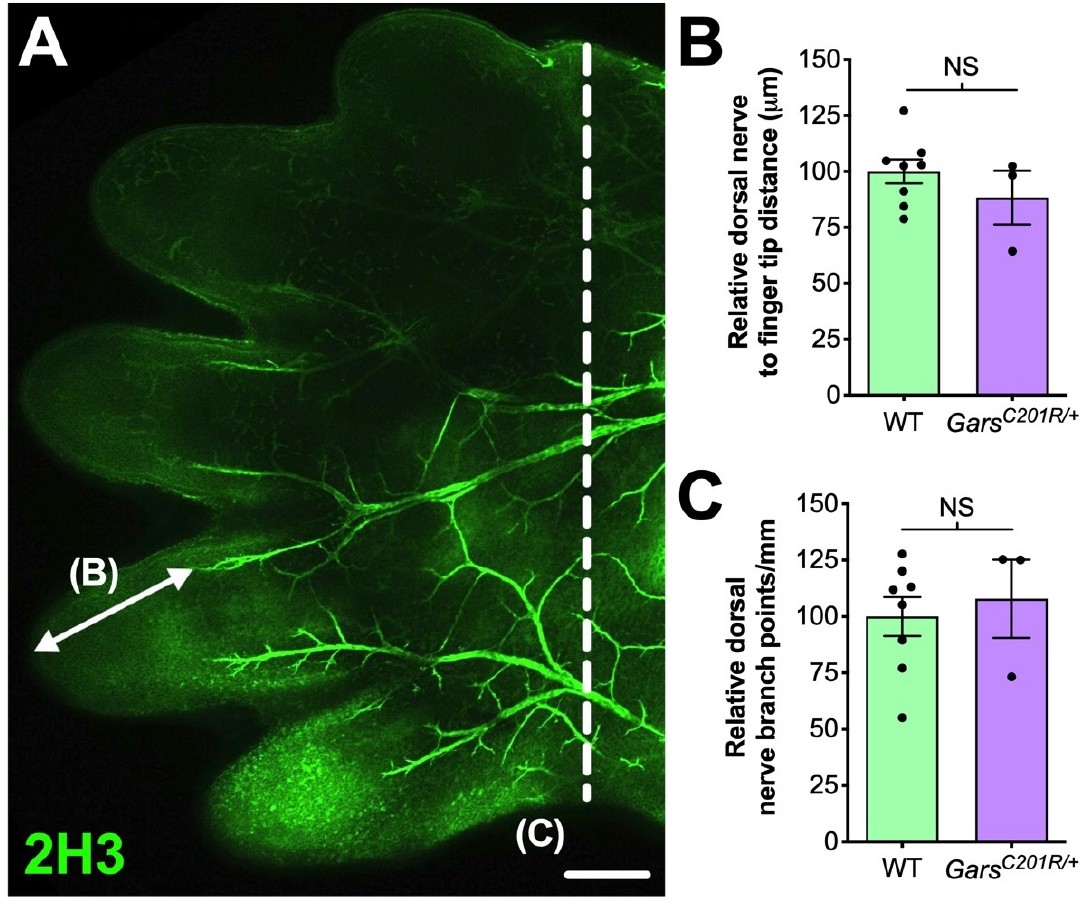
Sensory neurodevelopment appears normal in the forelimb of *Gars*^*C201R*/+^ embryos. (A) Representative single confocal plane, tile scan image of the dorsal aspect of an E13.5 wild-type forepaw stained for neurofilament (2H3, green). The arrow depicts distance from a major nerve branch ending to the tip of a finger, which was measured for B. The nerves to the left of the dashed line were used for branch analysis in C. Scale bar = 250 μm. (**B**, **C**) There was no difference between wild-type and *Gars*^*C201R*/+^ mice in sensory nerve extension into forepaw extremities (B, *P* = 0.328, unpaired *t*-test), nor in the amount of branching (C, *P* = 0.662, unpaired *t*-test). *n* = 3-8. *NS*, not significant; *WT*, wild-type.

### 3.2 Sensory populations are unaltered in a mouse model of ALS

We believe that the small, yet physiologically relevant, distortion of sensory populations is associated with aberrant mutant GlyRS-Trk receptor binding during development. To see whether it extends to other mouse models of neuromuscular disease, we analysed L1-L5 DRG from SOD1^G93A^ mice, an established model of amyotrophic lateral sclerosis (ALS), which displays a plethora of defects and dysfunctional pathways in peripheral, but mainly motor, nerves (Kim *et al.*, 2015; Nardo *et al.*, 2016). Lumbar DRG were dissected and immunohistochemically processed from SOD1^G93A^ and littermate control males at P30-31 (**Figure 3A**) and P100-101 (**Figure 3C**), representing pre-symptomatic and late disease stages, respectively. No distinctions in sensory populations were observed at either age (**Figure 3B** and **D**), indicating that the sensory subtype switch is not commonly observed in mouse models of neuromuscular diseases.

**Figure 3.**
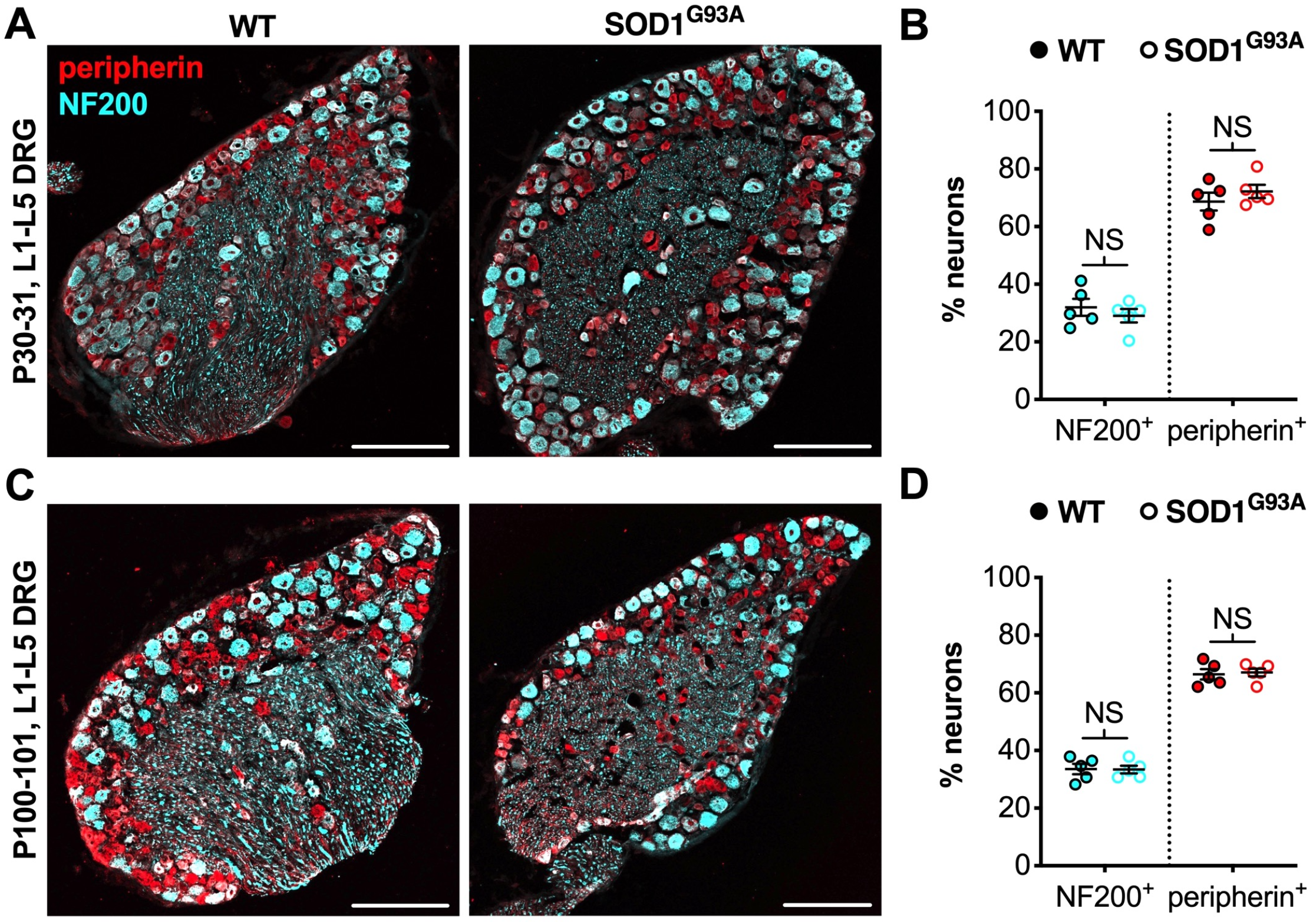
The CMT2D lumbar DRG sensory subtype switch is not observed in SOD1^G93A^ mice. (**A**, **C**) Representative immunofluorescence images of L1-L5 DRG sections from P30-31 (A) and P100-101 (C) wild-type and SOD1^G93A^ mice stained for NF200 (cyan) and peripherin (red). Scale bars = 200 μm. (**B**, **D**) There was no difference in the percentage of NF200^+^ and peripherin^+^ neurons between wild-type and SOD1^G93A^ mice at P30-31 (NF200 *P* = 0.460, peripherin *P* = 0.391; unpaired *t*-tests) nor P100-101 (NF200 *P* = 0.938, peripherin *P* = 0.782; unpaired *t*-tests). *n* = 5. *NS*, not significant; *WT*, wild-type. See also **Supplementary Figure S2**.

SOD1^G93A^ and *Gars*^*C201R*/+^ mice are maintained on different genetic backgrounds, and it appeared as though there may be a small difference in neuron populations between wild-types of the two strains. We therefore compared neuron proportions at one and three months in lumbar DRG from wild-type mice on a mixed C57BL/6 SJL background (SOD1^G93A^ control) versus a pure C57BL/6J background (*Gars*^*C201R*/+^ control). We observed a small, but significant difference between strains in the percentage of NF200^+^, but not peripherin^+^, neurons at one month, but not three months (**Supplementary Figure S2B** and **S2C**).

### 3.3 Long-range signalling endosome transport is unaffected in CMT2D sensory neurons

Axonal transport is reliant upon motor proteins traversing microtubule networks to deliver diverse cargoes from one end of an axon to the other (Guedes-Dias and Holzbaur, 2019). Anterograde transport from cell body to axonal terminal is key for delivering organelles, proteins and RNAs towards peripheral synapses. Connecting the axon tip to cell body, retrograde transport is needed for long-range delivery of autophagosomes and survival-promoting neurotrophic factors. Pre-symptomatic disturbances in axonal trafficking are thought to underlie, or at least contribute to, several neurological diseases (Sleigh *et al.*, 2019). Indeed, primary DRG neurons cultured from 12 day old *Gars*^*Nmf249*/+^ mice display reduced retrograde transport speeds of nerve growth factor (NGF)-loaded endosomes (Mo *et al.*, 2018), while reduced mitochondrial motility was also identified in sensory processes of 12 month old *Gars*^*C201R*/+^ mice (Benoy *et al.*, 2018). Disruption of two different cargoes suggests a broad transport impairment (e.g. due to microtubule dysfunction); however, *Gars*^*C201R*/+^ neurons were cultured from late symptomatic mice, thus the defective trafficking observed in this mutant may simply be a secondary consequence of neuropathology.

To analyse *Gars*^*C201R*/+^ transport in early symptomatic sensory neurons, we cultured primary thoracic and lumbar DRG neurons from one and three month old mice and assessed retrograde signalling endosome trafficking. These spinal levels were combined to obtain sufficient cell numbers for the assay (**Figure 1B**), and the timepoints were chosen to allow comparison with several other phenotypes assessed previously in this model (Sleigh *et al.*, 2014b, 2017a, 2017b). Cultures were incubated with fluorescently labelled atoxic binding fragment of tetanus neurotoxin (H_C_T-647), which is taken up by neurons and loaded into signalling endosomes containing Trk receptors and p75 neurotrophin receptor (p75NTR), when applied to media (Deinhardt *et al.*, 2006, 2007). Time-lapse confocal microscopy was performed to enable tracking of individual endosome kinetics (**Figure 4A**). Sensory neurons cultured from DRG do not always display a clearly visible axon initial segment and may bear several morphologically indistinguishable axon-like extensions/processes (Nascimento *et al.*, 2018), thus the trafficking analysed may not always be “axonal” transport *per se*.

**Figure 4.**
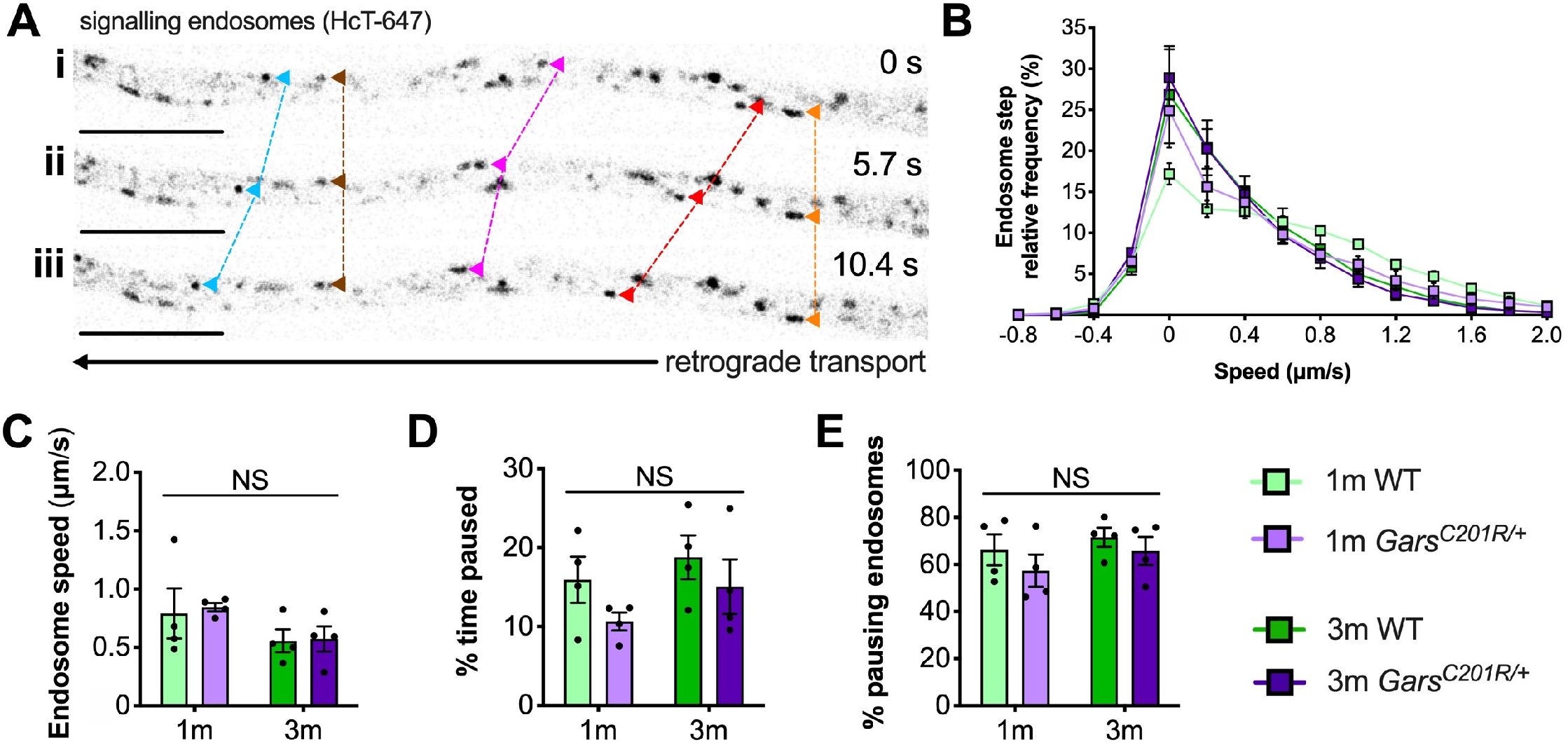
Axonal transport of signalling endosomes is unaffected in large area *Gars*^*C201R*/+^ sensory neurons. (**A**) Individual signalling endosomes loaded with fluorescently labelled H_C_T (H_C_T-647, grayscale inverted) were tracked and analysed from primary sensory neurons. Image series (*i-iii*) depicts retrograde (right to left) trafficking of distinct endosomes (arrowheads linked by dashed lines across time series) over time. The imaged neuron was cultured from a P97 wild-type female. Scale bars = 10 μm. (**B**) Frame-to-frame speed distribution curves indicate that there was no clear distinction in endosome kinetics between genotypes or timepoints. (**C-E**) This was confirmed by analysing average endosome speeds (C, genotype *P* = 0.792, age *P* = 0.076, interaction *P* = 0.884; two-way ANOVA), the percentage of time endosomes were paused for (D, genotype *P* = 0.123, age *P* = 0.206, interaction *P* = 0.779; two-way ANOVA), and the percentage of pausing endosomes (E, genotype *P* = 0.245, age *P* = 0.273, interaction *P* = 0.805; two-way ANOVA). *n* = 4. *1m*, 1 month; *3m*, 3 months; *NS*, not significant; *WT*, wild-type. See also **Supplementary Figure S3** and **S4**.

Contrary to the previous studies, overlapping histograms of endosome frame-to-frame speeds suggest that there is little difference in endosome dynamics between genotypes and timepoints (**Figure 4B**). This was confirmed by analysing average endosome speeds (**Figure 4C**), the percentage of time that endosomes were paused (**Figure 4D**), and the percentage of pausing endosomes (**Figure 4E**) per animal. There was also no difference in transport parameters between wild-type and *Gars*^*C201R*/+^ when axons were used as the experimental unit (**Supplementary Figure S3**); however, irrespective of the genotype, older cultures did display a general slowing of endosome speeds linked to increased pausing when compared in this manner.

Transport was assessed in larger area neurons only, because there is less frequent overlap of moving endosomes in wider processes, likely due to lower microtubule density (Ochs *et al.*, 1978), permitting greater tracking accuracy. Although not confirmed immunohistochemically, analysed cells were therefore very likely to be medium-large NF200^+^ sensory neurons (*i.e.* mechanosensitive or proprioceptive). Nonetheless, given the distortion in sensory subtypes in CMT2D lumbar DRG (Sleigh *et al.*, 2017a), it is possible that different neuron populations were analysed between genotypes. Thus, we measured cell body areas and process diameters from the neurons in which endosome transport was assessed (**Supplementary Figure S4**). There were no differences in these morphological properties, indicating that similar neurons were analysed across genotypes and timepoints.

### 3.4 Elevated GlyRS levels in select neurons are linked to the sensory subtype switch

GlyRS protein levels have previously been reported to be elevated in both *Gars*^*C201R*/+^ and *Gars*^*Nmf249*/+^ brains (Achilli *et al.*, 2009; Stum *et al.*, 2011), perhaps as a compensatory response to impaired protein function. To determine whether GlyRS levels are also altered in sensory neurons, we extracted and performed western blotting on C4-C8 and L1-L5 DRG from one month old *Gars*^*C201R*/+^ mice (**Figure 5A**). There was no difference in GlyRS levels in cervical ganglia; however, GlyRS was upregulated more than 2-fold in L1-L5 DRG (**Figure 5B**). This was corroborated by GlyRS immunofluorescence analysis in C4-C8 (**Figure 5C**) and L1-L5 (**Figure 5D**) ganglia. We also assessed L1-L5 DRG of one month old *Gars*^*Nmf249*/+^ mice and saw the same pattern of enhanced GlyRS fluorescence in a subset of DRG neurons (**Figure 5E**), thus indicating that the increase of mutant GlyRS levels in L1-L5 DRG is an early event in CMT2D pathogenesis.

**Figure 5.**
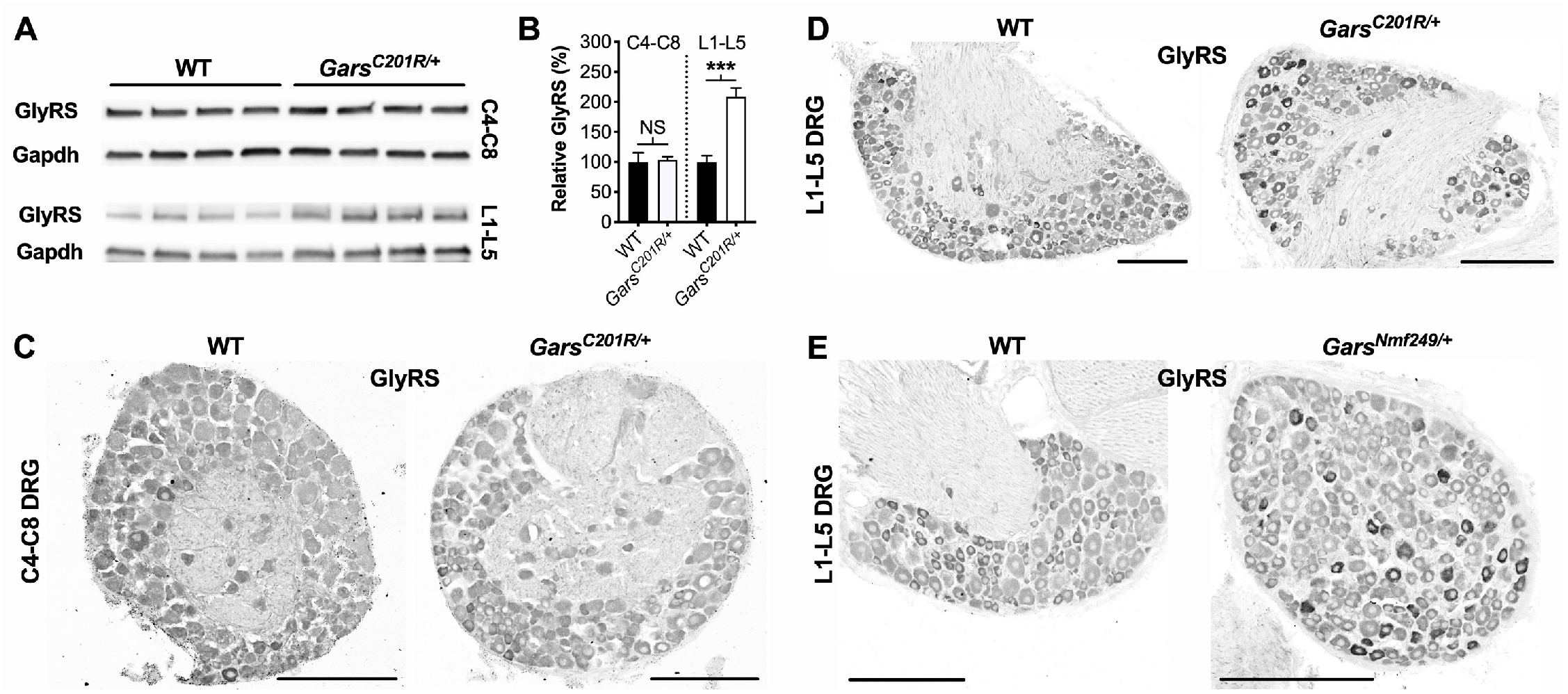
GlyRS protein levels are elevated in lumbar, but not cervical, DRG of CMT2D mice. (**A**) Representative western blot of C4-C8 (top) and L1-L5 (bottom) DRG lysates from one month old wild-type and *Gars*^*C201R*/+^ mice probed for GlyRS and the loading control Gapdh. (**B**) There was no difference between genotypes in GlyRS levels in cervical ganglia (*P* = 0.825; unpaired *t*-test; *n* = 5); however, GlyRS was elevated in L1-L5 DRG (*** *P* < 0.001, *NS* not significant; unpaired *t*-test; *n* = 4). (**C**, **D**) This was confirmed by immunofluorescence analysis of GlyRS in cervical (C) and lumbar (D) DRG sections from one month old wild-type and *Gars*^*C201R*/+^ mice. *n* = 4. GlyRS levels appear higher in many individual neurons with larger cell bodies in the mutant lumbar DRG (top right image). (**E**) These findings were replicated when comparing sections of L1-L5 ganglia from one month old wild-type and *Gars*^*Nmf249*/+^ mice. *n* = 3. Scale bars = 200 μm. *WT*, wild-type. See also **Supplementary Figure S5**.

Upon closer inspection, GlyRS immunofluorescence is marginally higher in some of the smaller area neurons of wild-type lumbar DRG; however, in both *Gars* mutants, the upregulation appears to be in larger neurons. To better characterise this, we co-stained *Gars*^*Nmf249*/+^ lumbar DRG for GlyRS and NF200 (**Figure 6A**). We found that ≈87% of neurons with increased GlyRS levels were also NF200^+^ (**Figure 6B**), which is a particularly high proportion considering that these mutant ganglia consist of only ≈22% NF200^+^ neurons (Sleigh *et al.*, 2017a). This suggests that there is a preferential increase of GlyRS in mechanosensitive and proprioceptive neurons. We then quantified the percentage of NF200^+^ neurons that showed elevated GlyRS and found that ≈32% showed the phenotype (**Figure 6B**), indicating that GlyRS is differentially upregulated even within this neuronal population, perhaps in a subgroup with a particular function.

**Figure 6.**
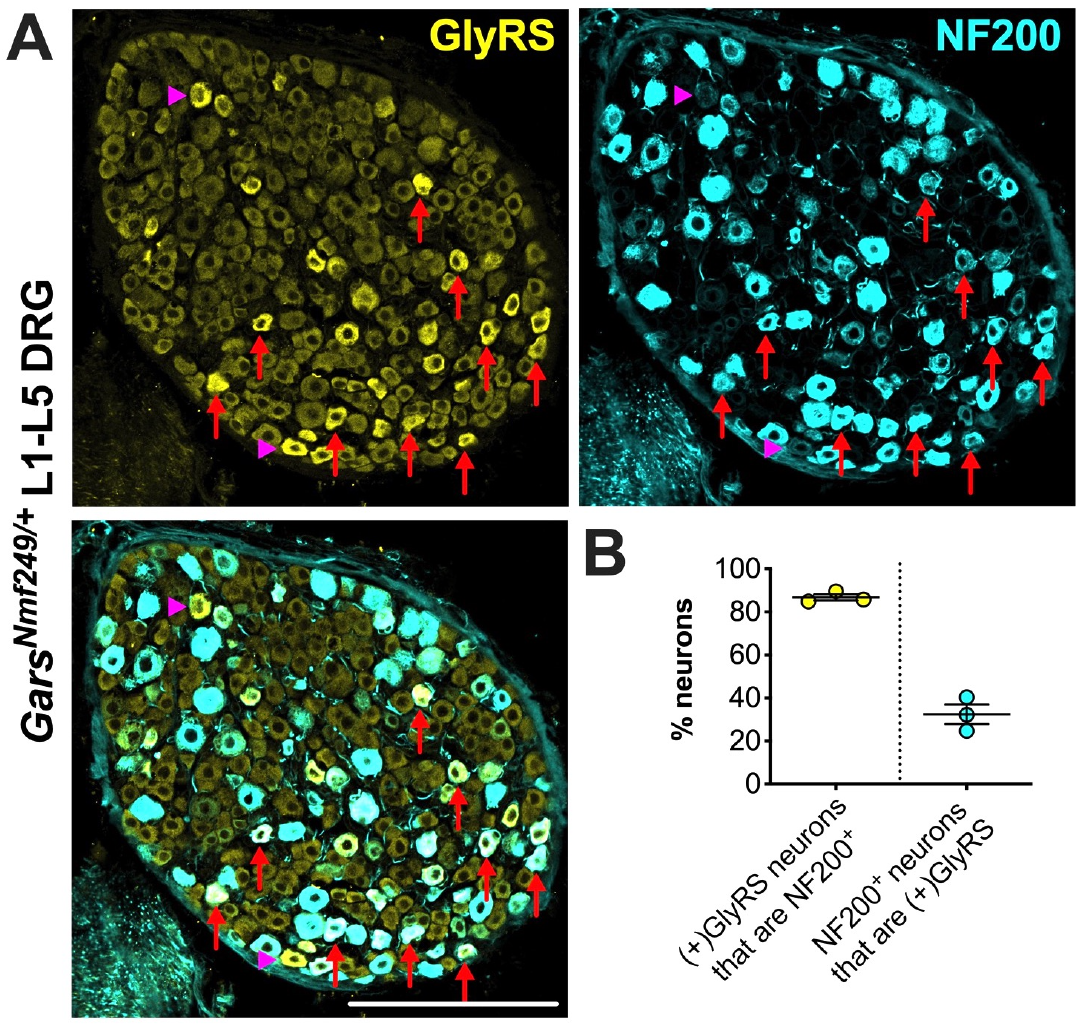
GlyRS is preferentially increased in some, but not all, NF200^+^ sensory neurons of *Gars^Nm249/+^* mice. (**A**) Representative immunofluorescence image of a L1-L5 DRG section from a one month old *Gars^Nm249/+^* mouse stained for GlyRS (yellow) and NF200 (cyan). Red arrows and magenta arrowheads highlight N200^+^ and NF200^−^ neurons, respectively, in which GlyRS was increased. Scale bar = 200 μm. (**B**) The majority of neurons in which GlyRS was upregulated, designated “(+)GlyRS”, express NF200; however, not all NF200^+^ cells have increased GlyRS levels. *n* = 3.

A global increase in ARS proteins would perhaps suggest dysfunction in a cellular process linked to aminoacylation, for instance protein translation, which is impaired a gain-of-function manner in CMT2D fly models (Niehues *et al.*, 2015). We therefore assessed whether upregulation is GlyRS-specific by analysing lumbar DRG levels of Kars-encoded lysyl-tRNA synthetase (LysRS) and Yars-encoded tyrosyl-tRNA synthetase (TyrRS) (**Supplementary Figure S5A**). LysRS was chosen because, with GlyRS, it is the only other dual-localised synthetase functioning in both cytoplasm and mitochondria, while TyrRS was chosen because there is strong evidence that mutations in its encoding gene, YARS1, cause CMT (Wei *et al.*, 2019). *Gars*^*C201R*/+^ L1-L5 DRG show no change in LysRS levels and only a small, but significant, rise in TyrRS (**Supplementary Figure S5B**). We confirmed the lack of LysRS upregulation by staining lumbar ganglia from *Gars*^*Nmf249*/+^ mice (**Supplementary Figure S5C**). TyrRS immunohistochemistry was attempted, but the resulting staining pattern was not consistent with a cytoplasmic tRNA synthetase.

Together, these data indicate that there is a selective increase in *Gars* expression that occurs predominantly in NF200^+^ cells and only at spinal levels displaying a developmental perturbation of sensory neuron fate.

## 4. Discussion

### 4.1 Sensory phenotypes are restricted to the lower limbs of CMT2D mice

CMT2D mice display developmental phenotypes in sensory neurons innervating the hindpaws (Sleigh *et al.*, 2017a). To extend these analyses and determine whether the upper limb predominance of patients is replicated, we assessed afferent nerves from cervical spinal levels in *Gars*^*C201R*/+^ mice. The lumbar phenotypes of subtype switching and impaired axon branching were not present in sensory neurons targeting forepaws (**Figure 1**, **2** and **Supplementary Figure 1**), suggesting that anatomical location has a considerable bearing on neurodevelopmental pathology. Given that these phenotypes are developmental and do not progress in severity from P1 to three months, it is unlikely that they occur in cervical DRG later in disease.

Although mutations in *GARS1* frequently cause hands to be affected before and more severely than feet, there are examples where the opposite occurs (Sivakumar *et al.*, 2005; Forrester *et al.*, 2020); albeit it is unclear as to whether this also applies to sensory symptoms. In a similar manner, the *Gars^C201R^* mutation could therefore simply preferentially impact lower limbs. However, restriction of weakness to CMT2D patient feet is rare. So, what could be driving the differential pathology between lumbar and cervical ganglia in mice? It may be due to a distinction in the proportions of sensory subtypes. In wild-type mice, NF200 is expressed by ≈41% of neurons in L1-L5 DRG (Sleigh *et al.*, 2017a), whereas ≈56% are N200+ in cervical ganglia (**Supplementary Figure S2A**). If mutant GlyRS aberrantly interacts with Trk receptors pre-natally, thus impacting sensory development and skewing the proportions of functional subclasses (Sleigh *et al.*, 2017a), then GlyRS is likely to have less impact on ganglia that have a more equal balance between NF200^+^ and peripherin^+^ cells, as is observed in C4-C8 DRG. Alternatively, spinal level distinctions may be caused by differences in the amount or kinetics of GlyRS secretion or Trk expression. DRG at different spinal levels develop asynchronously (Lawson and Biscoe, 1979) and possess divergent transcription factor profiles (Lai *et al.*, 2016), which may also contribute to the lower limb predominance observed in CMT2D mice. Indeed, the transcription factors neurogenin 1 and 2, which drive two distinct waves of neurogenesis required for segregation of major classes of Trk-expressing sensory neurons, are differentially required by cervical and more caudal sensory ganglia (Ma *et al.*, 1999).

Irrespective of the cause, experiments presented here highlight the importance of comparative anatomy in mouse models of neuromuscular disorders to enhance understanding of pathomechanisms. It remains to be seen whether such variations are also observed in the motor nervous system of CMT2D mice, although differential susceptibility of muscles to NMJ denervation has been previously reported (Seburn *et al.*, 2006; Sleigh *et al.*, 2014b; Spaulding *et al.*, 2016).

### 4.2 Developmental perturbation of sensory fate is not a common phenotype

A difference in sensory neuron populations has also been reported in lumbar DRG of mice modelling spinal muscular atrophy (SMA) (Shorrock *et al.*, 2018). We therefore aimed to determine whether this phenotype is a general feature of mouse models of neuromuscular disease, as this would cast doubt on the aberrant binding of mutant GlyRS to Trk receptors as being the cause of impaired sensory development in mutant *Gars* mice. SOD1^G93A^ mice modelling ALS show a variety of defects in sensory neurons (Sassone *et al.*, 2016; Vaughan *et al.*, 2018; Seki *et al.*, 2019); however, they do not display a subtype switch in lumbar DRG (**Figure 3**). Moreover, we recently found that mice modelling a developmental form of SMA caused by loss-of-function mutations in BICD2 also do not show this phenotype (Rossor *et al.*, 2020). Together, these data confirm that perturbed sensory development is not commonly observed in mouse models of neurodegeneration.

We did however see a small difference in lumbar DRG between wild-type littermates of SOD1^G93A^ and *Gars*^*C201R*/+^ mice (**Supplementary Figure S2B** and **S2C**), which are maintained on different genetic backgrounds, suggesting that genetic background may subtly influence sensory neuron populations, likely contributing to previously reported disparities in sensation between strains (Crawley *et al.*, 1997). This result is not overly surprising considering that mouse genetic background can influence even gross anatomical features such as the number of spinal levels (Rigaud *et al.*, 2008).

### 4.3 Long-range transport is not universally impaired in CMT2D sensory neurons

Deficits in axonal transport contribute to many different genetic neuropathies (Prior *et al.*, 2017; Beijer *et al.*, 2019), and its early involvement in disease may be a common driver in peripheral nerve selectivity typical of many forms of CMT2. Indeed, disruption of long-range trafficking has been identified in sensory neurons cultured from CMT2D DRG (Benoy *et al.*, 2018; Mo *et al.*, 2018). Contrastingly, we found no difference in retrograde transport of signalling endosomes between wild-type and *Gars*^*C201R*/+^ at one and three months of age (**Figure 4** and **Supplementary Figure S3**). Nonetheless, if thoracic DRG show limited to no pathology, by combining thoracic DRG with L1-L5 ganglia we may have masked a lumbar-specific DRG transport phenotype. Additionally, the medium and associated supplements in which DRG neurons were cultured vary across studies. Neuronal activity can impact the rate and quantity of axonal transport (Sajic *et al.*, 2013; Wang *et al.*, 2016), while proteins such as neurotrophic factors, which are present in primary neuron media, can affect neuronal activity (Dombert *et al.*, 2017). Though unlikely, it is possible therefore that the medium in which our neurons were grown may have selectively enhanced *Gars*^*C201R*/+^ transport masking a trafficking deficiency.

Mo *et al.* (2018) identified a slow-down in NGF-containing endosomes tracked in sensory neurons cultured from 12 day old *Gars*^*Nmf249*/+^ mice. Firstly, if transport disruption correlates with overall disease burden, then the more severe mutant allele is more likely to display a defect than the milder *Gars*^*C201R*/+^ mutant. Equally as importantly, the assayed neurons in the two studies were perhaps different. NGF binds to TrkA, which is expressed by nociceptors (**Supplementary Figure S1A**), thus transport was probably assessed in noxious stimulus-sensing peripherin^+^ neurons from *Gars*^*Nmf249*/+^ mice. The atoxic binding fragment of tetanus neurotoxin that we used to assess transport is taken up into multiple populations of signalling endosomes (Villarroel-Campos *et al.*, 2018); however, we focused our analyses on large area neurons with wide processes (**Supplementary Figure S4**), likely to be either TrkB+ mechanoreceptors or TrkC+ proprioceptors.

Benoy *et al.* (2018) identified that *in vitro* sensory neurons cultured from 12 month old *Gars*^*C201R*/+^ mice had an almost complete impairment in mitochondrial motility. The disparity with this study could simply reflect the analysed cargo, *i.e.* mitochondrial, but not endosomal, transport is altered in this model. Alternatively, the difference may be due to the age at which the neurons were tested (one and three months versus 12 months). Supporting this idea, we have previously shown that *Gars*^*C201R*/+^ sensory neurons cultured from one month old animals show normal neurite/process outgrowth (Sleigh *et al.*, 2017a); however, this was defective in 12 month old cells (Benoy *et al.*, 2018). Decreased neuronal health may therefore be contributing to reduced mitochondrial motility, as may the process of aging. Indeed, we have previously reported that the dynamics of signalling endosome transport *in vivo* remain unaltered in aged wild-type mice (Sleigh and Schiavo, 2016; Sleigh *et al.*, 2020), whereas mitochondrial transport is known to be altered in old animals (Mattedi and Vagnoni, 2019).

Our long-rang retrograde transport data indicate that cargo trafficking is not globally disrupted in all CMT2D sensory neurons during early disease stages. To unravel the significance of axonal transport impairments to CMT2D aetiology, it will be important to assess trafficking of a variety of different cargoes both in sensory and motor neurons of mutant *Gars* models. Given the complexity of the *in vivo* environment, the kinetics of axonal transport are not always replicated *in vitro* (Sleigh *et al.*, 2017c), hence analysis of mutant *Gars* mouse transport should also be extended to peripheral nerves *in vivo* (Gibbs *et al.*, 2016, 2018).

### 4.4 GlyRS elevation is not a simple compensatory mechanism

Neuropathy-causing *GARS1* mutations differentially impact the enzymatic activity, with some fully ablating it, whilst others having little effect (Oprescu *et al.*, 2017). The charging function of GlyRSC201R and GlyRSP278KY in *Gars*^*C201R*/+^ and *Gars*^*Nmf249*/+^ mice, respectively, were originally reported as unaffected (Seburn *et al.*, 2006; Achilli *et al.*, 2009; Stum *et al.*, 2011). However, a re-evaluation under Michaelis-Menten kinetic conditions suggests that GlyRSP278KY has severely decreased kinetics and cannot support yeast viability, commensurate with a loss-of-function (Morelli *et al.*, 2019). Moreover, GlyRSC201R aminoacylation was analysed indirectly in brain lysates that had a 3.8-fold increase in GlyRS (Achilli *et al.*, 2009), which could mask a charging deficiency. Accordingly, brains from severe homozygous *Gars^C201R/C201R^* mice showed a 60% decrease in aminoacylation despite an 8.2-fold increase in GlyRS. Further supporting a GlyRSC201R loss-of-function, wild-type *GARS1* overexpression in sub-viable homozygous *Gars^C201R/C201R^* animals is able to restore post-natal viability (Motley *et al.*, 2011). Similar to *Gars*^*C201R*/+^ brains, GlyRS levels were reported to be higher in *Gars*^*Nmf249*/+^ cerebellum, although this was not quantified (Stum *et al.*, 2011). It is therefore possible that GlyRS levels are elevated in CMT2D tissues as a compensatory response to diminished aminoacylation.

To test this hypothesis in sensory tissue, we analysed GlyRS protein in CMT2D DRG (**Figure 5**). Coinciding with the perturbation of sensory neuron fate, we observed enhanced GlyRS levels in lumbar, but not cervical, ganglia of mutant *Gars* mice. Furthermore, the increase was not observed in all lumbar sensory neurons, but preferentially in a portion of NF200^+^ neurons (**Figure 6**). This argues against GlyRS upregulation being a compensatory response to impaired charging, an alteration in a non-canonical function (Johanson *et al.*, 2003; Park *et al.*, 2012; Mo *et al.*, 2016), or that mutant GlyRS protein stability is altered, because if any of those scenarios were true, then GlyRS increase would also likely occur in cervical DRG and across all sensory neurons equally. That is, unless there is a greater requirement for glycine charging in cell bodies of larger sensory neurons with the longest axons (*i.e.* those innervating lower, but not upper, limbs). Contradictory to this idea, GlyRS levels were enhanced in only about a third of NF200^+^ neurons in mutant lumbar DRG, suggesting that a particular subset may be selectively impacted by disease.

To further tease apart the basis for increased *Gars* expression, we assessed levels of additional ARS proteins. We found that LysRS remained unchanged and there was only a small increase in TyrRS in CMT2D DRG (**Supplementary Figure S5**), indicating that there is no global increase in tRNA synthetase in response to GlyRSC201R expression. We therefore observe a GlyRS-specific upregulation, preferentially occurring in a subdivision of NF200^+^ neurons and only in ganglia that display neuropathology. Why might this be the case? Neuropathy-associated *GARS1* mutations have been shown to impair GlyRS localisation in neuron-like cell lines (Antonellis *et al.*, 2006; Nangle *et al.*, 2007), which could cause build-up in the soma, although, once again, if this were the cause of increased GlyRS levels then it would probably not be so selectively upregulated. The GlyRS elevation is only present in DRG that display a developmental perturbation in sensory neuron fate, suggesting that the two phenotypes may be linked. Perhaps the NF200^+^ neurons resident in lumbar ganglia are under stresses not experienced by neighbouring subtypes. The integrated stress response (ISR), which is linked to amino acid deprivation (Pakos-Zebrucka *et al.*, 2016), may be especially activated in these cells. Consistent with impaired protein translation reported in CMT2D fly models (Niehues *et al.*, 2015), the ISR causes a global downregulation of cap-dependent translation of mRNAs, except for a select few that possess upstream open reading frames (uORFs) in their 5’-UTRs, which under non-stressed conditions usually restrict translation initiation of the main downstream ORF (Barbosa *et al.*, 2013). Although not classically thought of as an ISR-associated gene, human and mouse *GARS1* express two mRNA isoforms, one of which possesses an uORF that may, under conditions of stress, play a role in the observed GlyRS increase (Alexandrova *et al.*, 2015). However, some KARS1 variants also possess an uORF (AceView, NCBI) and LysRS levels remained unchanged. Alternatively, the increase may be an active, compensatory response by a subset of NF200^+^ cells to combat degeneration. Indeed, the NF200^+^ class of neurons includes vibration-sensing mechanoreceptors, which are most impacted in CMT2D patients (Sivakumar *et al.*, 2005).

### 4.5 Conclusion

Sensory dysfunction of *GARS1*-neuropathy patients and mouse models of CMT2D is chronically understudied. This is unsurprising given the relative severity of motor symptoms; however, by studying pathology in both types of peripheral nerve and performing comparative anatomical studies on mouse motor and sensory nervous systems, we are much more likely to determine key pathomechanisms causing the selective pathology characteristic of CMT. Here, we have made four key discoveries: 1) sensory pathology is not equal across all CMT2D ganglia, thus anatomical location dictates disease involvement; 2) perturbed sensory neuron fate is not a general feature of different neuromuscular disease models, supporting its specificity to *GARS1* neuropathy; 3) signalling endosome trafficking in a sub-population of *Gars*^*C201R*/+^ sensory neurons remains unaffected, indicating that a widespread CMT2D defect in axonal transport is unlikely; and 4) *Gars* expression is selectively enhanced in NF200^+^ lumbar DRG neurons and is thus linked to the subtype switch, perhaps in response to active degeneration.

## Acknowledgements

The authors would like to thank Drs. Emily Spaulding and Robert Burgess (The Jackson Laboratory, Bar Harbor, ME) for providing *Gars*^*Nmf249*/+^ tissue and commenting on the manuscript, James Dick (University College London) for genotyping the P100-101 SOD^G93A^ and wild-type mice, and Drs. Alexander M. Rossor and David Villarroel-Campos (University College London) for critical comments on the manuscript.

## Funding

This work was supported by the Medical Research Council Career Development Award (MR/S006990/1) [JNS], the Wellcome Trust Sir Henry Wellcome Postdoctoral Fellowship (103191/Z/13/Z) [JNS], the Wellcome Trust Senior Investigator Award (107116/Z/15/Z) [GS], the European Union’s Horizon 2020 Research and Innovation programme under grant agreement 739572 [GS], and the UK Dementia Research Institute Foundation award (UKDRI-1005) [GS].

## Author Contributions

JS conceived the experiments; JS, AM, TA, YZ performed the research; JS analysed the data; GS provided expertise and discussion; JS wrote the manuscript. All authors approved submission of this work.

## Conflict of Interest

None declared.

## Supplementary Figures

**Supplementary Figure 1.**
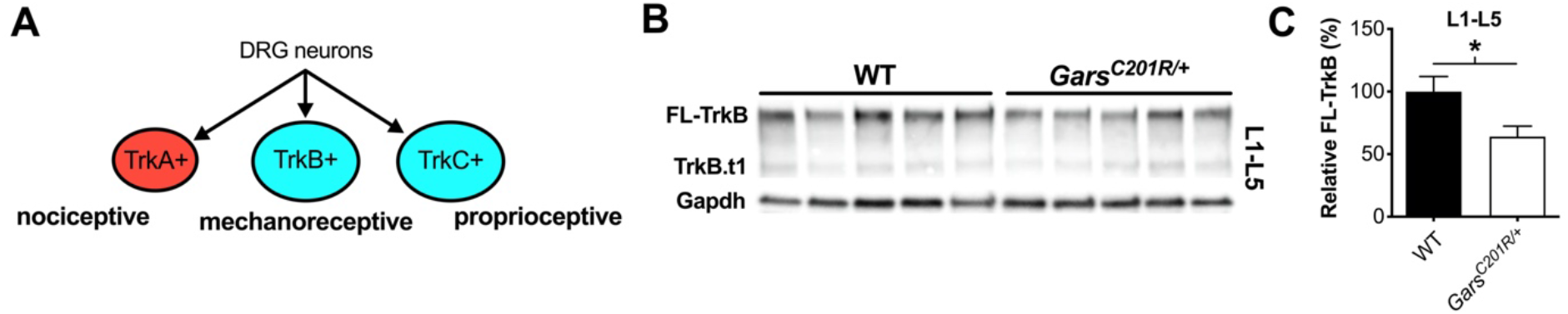
TrkB protein levels are lower in *Gars^C201R/+^*lumbar DRG consistent with fewer mechanosensory neurons. (**A**) Trk receptors A, B and C are expressed in post-natal DRG sensory neurons that function in nociception, mechanosensation and proprioception, respectively (Montaño et al., 2010). TrkA preferentially binds to NGF, TrkB to BDNF and NT-4, and TrkC to NT-3. (**B**) Representative western blot of L1-L5 DRG lysates from one month old wild-type and *Gars^C201R/+^* mice probed for TrkB and the loading control Gapdh. *FL-TrkB*, full-length TrkB; *TrkB.t1*, truncated TrkB. (**C**) Densitometry analysis indicates that CMT2D lumbar DRG have lower levels of FL-TrkB, likely reflecting the sensory subtype switch previously identified in lumbar DRG (Sleigh et al., 2017a). * *P* = 0.040, unpaired *t-*test. *n* = 5. *WT*, wild-type. Related to **Figure 1**.

**Supplementary Figure 2.**
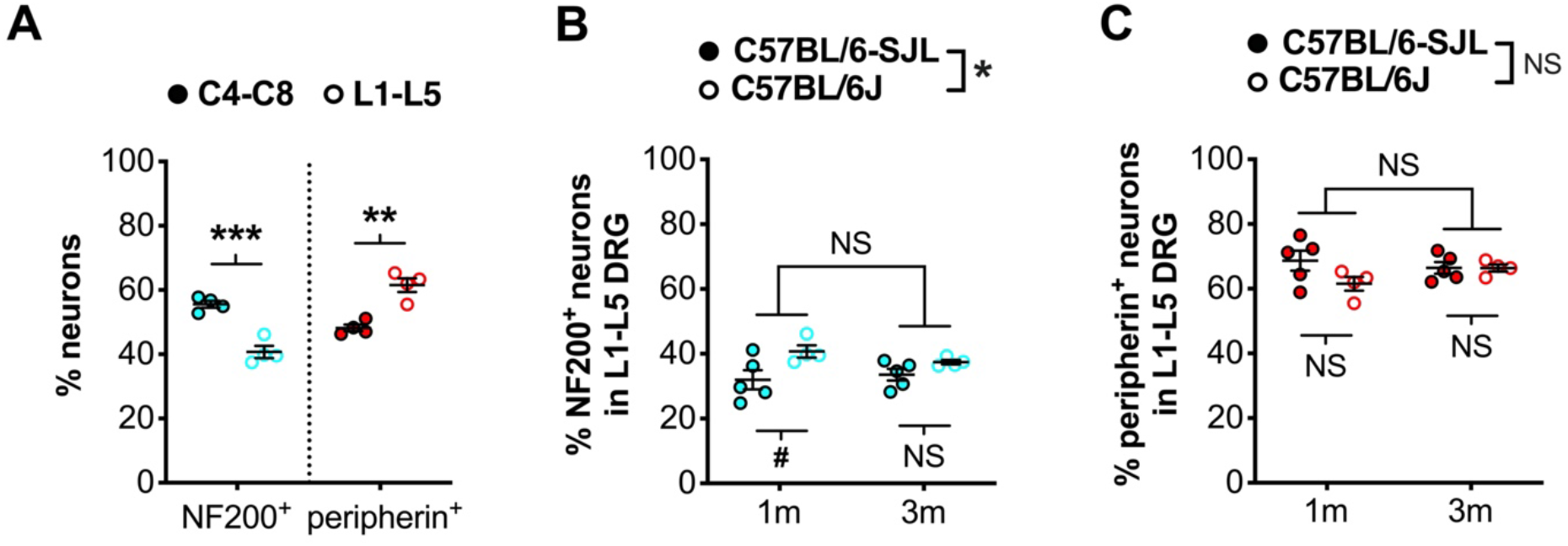
A large difference in sensory neuron populations is observed between cervical and lumbar ganglia. (**A**) C4-C8 DRG possess significantly more NF200+ neurons and significantly fewer peripherin+ cells than L1-L5 DRG. C4-C8 data are also presented as “*wild-type*” in **Figure 1D**. *** *P* < 0.001, ** *P* < 0.01; unpaired *t*-test. (**B**) There is a small difference in the percentage of NF200+ neurons in L1-L5 DRG between mice on a pure C57BL/6J background and a mixed C57BL/6-SJL background (strain * *P* = 0.011, age *P* = 0.694, interaction *P* = 0.277; two-way ANOVA). ^#^ *P* < 0.05, *NS* not significant; Sidak’s multiple comparisons test. (**C**) No difference is observed between strains in the percentage of peripherin+ neurons in lumbar ganglia (strain *P* = 0.140, age *P* = 0.576, interaction *P* = 0.144; two-way ANOVA). *NS* not significant; Sidak’s multiple comparisons test. The one month C57BL/6J data in B and C are presented as “*L1-L5*” in panel A. Data from C57BL/6J lumbar ganglia were generated for a previous study (Sleigh et al., 2017a). C57BL/6-SJL data are also presented as “*wild-type*” in **Figure 3B** and **D**. *N.b.*, statistically compared DRG datasets were independently stained, imaged and quantified, which may account for some variability. *n* = 4-5 *1m*, 1 month; *3m*, 3 months. Related to **Figure 1** and **3**.

**Supplementary Figure 3.**
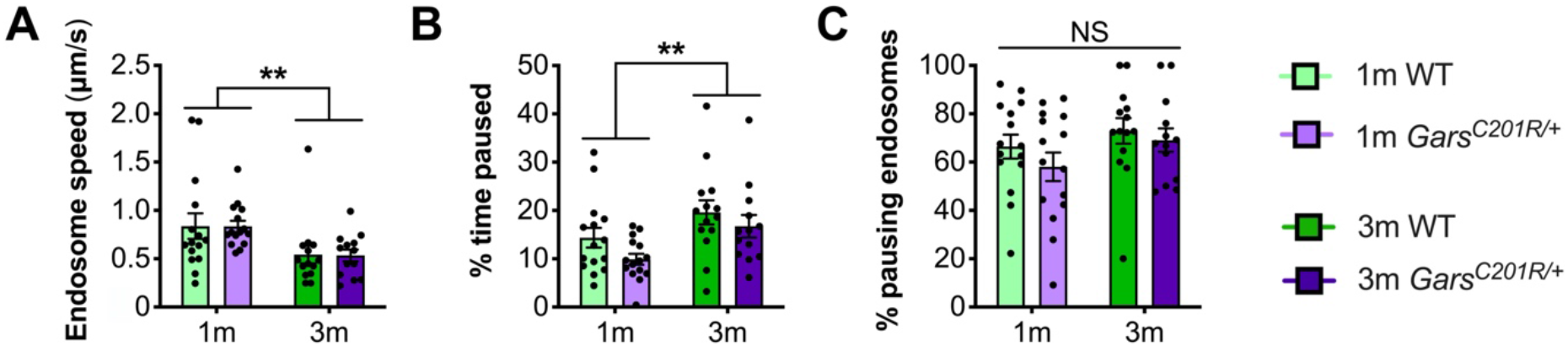
No difference in endosome transport between genotypes is found using neuronal process as the experimental unit. (**A-C**) A lack of distinction in endosome dynamics between genotypes was confirmed using processes as the experimental unit. There is no difference between wild-type and *Gars^C201R/+^* neurons in average endosome speeds (A, genotype *P* = 0.948, age ** *P* = 0.002, interaction *P* = 0.979; two-way ANOVA), percentage of time endosomes paused for (B, genotype *P* = 0.077, age ** *P* = 0.005, interaction *P* = 0.700; two-way ANOVA), or percentage of pausing endosomes (C, genotype *P* = 0.259, age *P* = 0.107, interaction *P* = 0.671; two-way ANOVA). However, there was a significant difference between timepoints in endosome speed (A) and % time paused (B), suggesting that there may be an age-related slow-down in endosome transport in cultured DRG sensory neurons. *n* = 13-15. *1m*, 1 month; *3m*, 3 months; *NS*, not significant; *WT*, wild-type. Related to **Figure 4**.

**Supplementary Figure 4.**
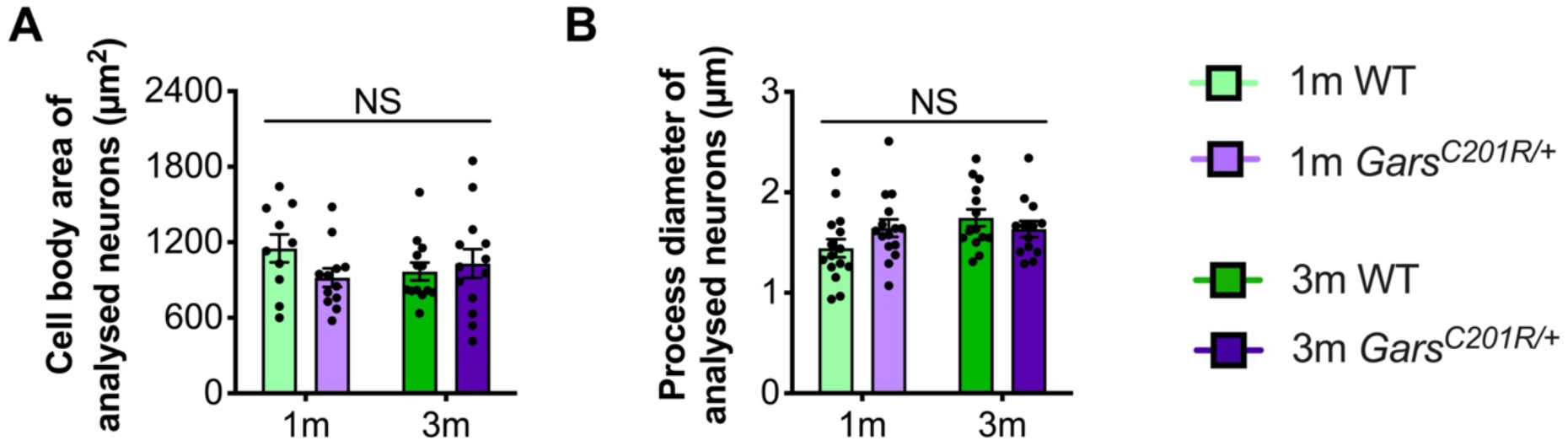
Large area sensory neurons of similar size between genotypes were assessed in the endosome transport assay. To ensure that retrograde transport of signalling endosomes was analysed in similar neuron subtypes between genotypes and across timepoints, the cell body area (**A**) and process diameter (**B**) of each tracked neuron were measured. No difference in either morphological property was detected, indicating that transport was assessed in comparable sensory neuron subtypes. *n* = 4, but with 10-15 individual neurons depicted. Two-way ANOVAs using either animal (*n* = 4) or process (*n* = 10-15) as the test replicate both indicated that there were no significance differences in age, genotype or their interaction. *N.b.,* at the one month timepoint, cell bodies were not measured in one experimental replicate, although their similar size was confirmed visually and their process diameters were no different from all other cultures (B). *1m*, 1 month; *3m*, 3 months; *NS*, not significant; *WT*, wild-type. Related to **Figure 4**.

**Supplementary Figure 5.**
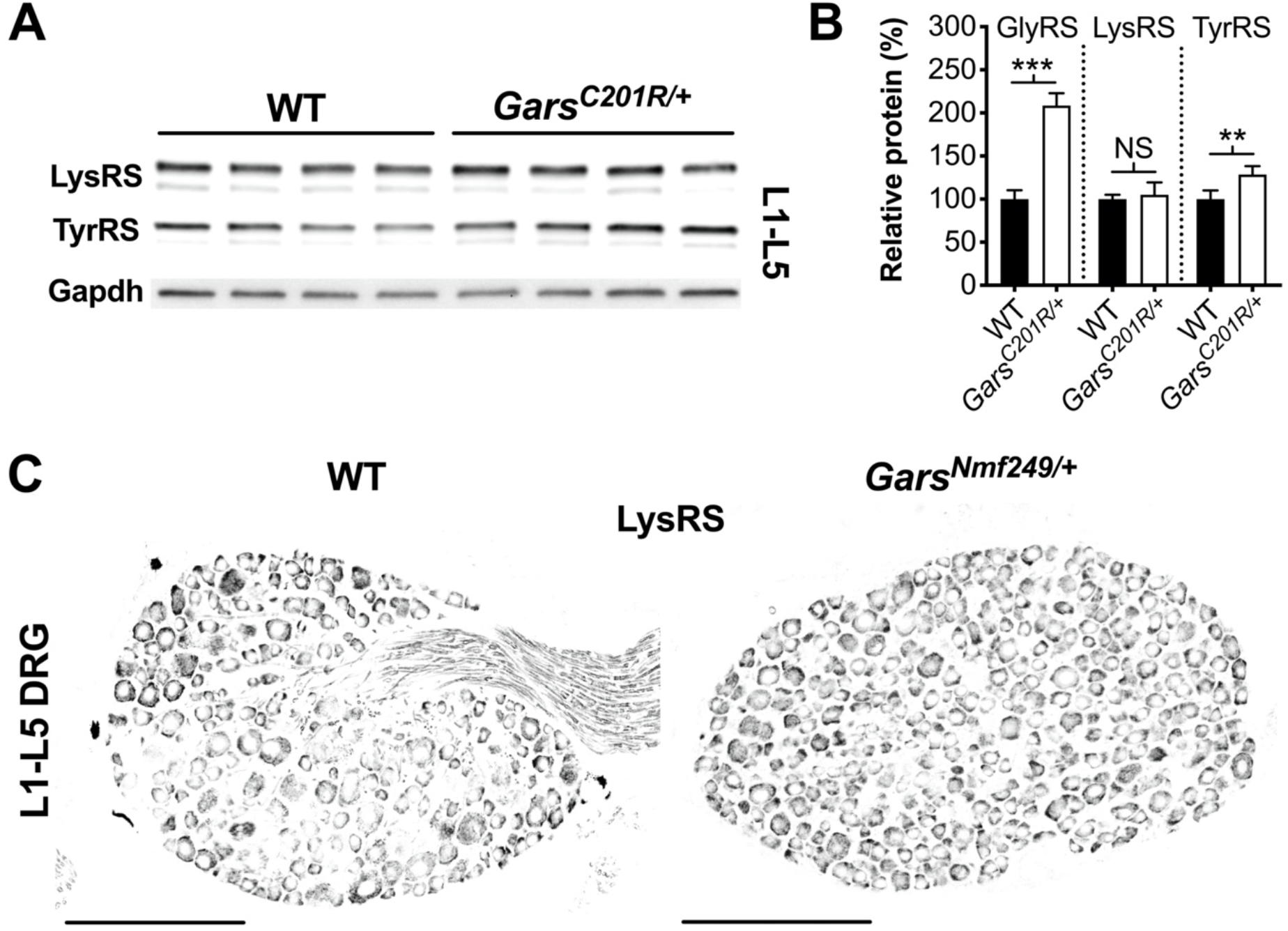
ARS proteins are not generally upregulated in lumbar DRG of CMT2D mice. (**A**) Representative western blot of L1-L5 DRG lysates from one month old wild-type and *Gars^C201R/+^* mice probed for aminoacyl-tRNA synthetase enzymes LysRS and TyrRS, and the loading control Gapdh. (**B**) Densitometry analysis indicates that LysRS levels did not differ between genotypes. There was a small, but significant, increase in TyrRS levels in *Gars^C201R/+^* lumbar DRG, but the change was much less than that observed for GlyRS (densitometry analysis from **Figure 5** included for comparison). *** *P* < 0.001, ** *P* < 0.01, NS, not significant; unpaired *t*-test. *n* = 4. (**C**) Representative immunofluorescence analysis of LysRS in L1-L5 ganglia sections from one month old wild-type and *Gars^Nmf249/+^* mice. Unlike GlyRS (**Figure 6**), but consistent with the *Gars^C201R/+^* LysRS western blot (A, B), individual neurons did not display an increase in LysRS levels. *n* = 3. *N.b.*, the positive LysRS signal observed in axons (wild-type image) was also seen in the secondary only control, and was therefore likely to be non-specific. Scale bars = 200 μm. *WT*, wild-type. Related to **Figure 5** and **6**.

## Notes

### Competing Interest Statement

The authors have declared no competing interest.

